# Neurocognitive mechanisms of d-cycloserine augmented single-session exposure therapy for anxiety

**DOI:** 10.1101/615757

**Authors:** Andrea Reinecke, Alecia Nickless, Michael Browning, Catherine J. Harmer

## Abstract

**Objective:** Drugs targeting the N-Methyl-D-aspartic acid (NMDA) system and the ability to learn new associations have been proposed as potential adjunct treatments to boost the success of exposure therapy for anxiety disorders. However, the effects of the NMDA partial agonist d-cycloserine on psychological treatment have been mixed. We investigated potential neurocognitive mechanisms underlying the clinical effects of d-cycloserine-augmented exposure, to inform the optimal combination of this and similar agents with psychological treatment.

**Methods:** Unmedicated patients with panic disorder were randomised to single-dose d-cycloserine (250mg; N=17) or matching placebo (N=16) 2hrs before one session of exposure therapy. Neurocognitive markers were assessed one day after treatment, including reaction-time based threat bias for fearful faces and amygdala response to threat. Clinical symptom severity was measured using self-report and clinician-rated scales the day before and after treatment, and at 1- and 6-months follow-up. Analysis was by intention-to-treat.

**Results:** One day after treatment, threat bias for fearful faces and amygdala threat response were attenuated in the drug compared to the placebo group. Lower amygdala magnitude predicted greater clinical improvement during follow-up across groups. D-cycloserine led to greater clinical recovery at 1-month follow-up (d-cycloserine 71% versus placebo 25%).

**Discussion:** D-cycloserine-augmented single-session exposure therapy reduces amygdala threat response, and this effect predicts later clinical response. These findings highlight a neurocognitive mechanism by which d-cycloserine may exert its augmentative effects on psychological treatment and bring forward a marker that may help understand and facilitate future development of adjunct treatments with CBT for anxiety disorders. (D-cycloserine Augmented CBT for Panic Disorder; clinicaltrials.gov; NCT01680107)

## Introduction

There has been increasing interest in the combination of exposure-based cognitive-behaviour therapy (CBT) for anxiety disorders with drugs targeting synaptic plasticity, to improve clinical outcome. The N-methyl-D-aspartate (NMDA) receptor partial agonist d-cycloserine has been shown to significantly facilitate clinical response to exposure under some conditions (1, 2), but the optimal methods for combination remain to be further identified and could be informed by characterisation of neurocognitive mechanisms underlying the clinical effects of intervention.

We have recently developed a single-dose CBT testbed that allows the assessment of early effects of treatment on such neurocognitive markers and their contribution to clinical improvement (3). Administering a single session of exposure therapy for panic disorder led to no clinical changes on the day after treatment, but resulted in a substantial decrease in threat bias for fearful faces. At 1-month follow-up and without additional interim treatment, clinical improvement was apparent, with 1/3 of treated patients fulfilling criteria for recovery. Importantly, clinical improvement was driven by the magnitude of threat bias measured on the day after treatment, with lower bias predicting improved clinical outcome. These findings suggest that a reduction of threat bias may be a key mechanism of action in exposure therapy, and they bring forward a single-dose CBT methodology that may help ascertain the mechanisms of action underlying pharmacological-psychological combination treatments. Such a focus on the mechanisms underpinning recovery with pharmacological exposure therapy can inform optimal combination of different treatment components, and it can provide an experimental medicine model for identification of novel agents.

This study aimed to characterise the effects of d-cycloserine with single-session CBT on neurocognitive markers important in panic disorder, including threat bias for fearful faces previously shown to mediate clinical effects of single-session CBT (3) and amygdala threat response previously shown to normalise after four sessions of exposure therapy in panic disorder (4). We hypothesised that the d-cycloserine group compared to placebo would show reduced threat bias and amygdala threat response after single-session CBT, and that the magnitude of these markers would predict clinical symptom changes during follow-up.

## Methods and Materials

### Participants

Thirty-three patients with a current panic disorder diagnosis and at least moderate agoraphobic avoidance were recruited from the community. Diagnosis was assessed using the Structured Clinical Interview for DSM-IV Axis I Disorders; avoidance was measured using the Structured Panic Assessment Interview (“yes” response to >2 situations listed under “(2) Avoidance”) (5, 6). Exclusion criteria were insufficient English skills, current or past psychotic or bipolar disorder, substance abuse/ dependence or epilepsy, current primary depressive disorder, CNS-active medication during the last 6 weeks, current medication with cycloserine, ethionamide or isoniazid, exposure-based CBT for panic during the last 3 months, pregnancy or lactation, and severe renal insufficiency or other serious medical conditions that may put the participant at risk. Occasional benzodiazepine or beta-blocker medication (15% of participants) was not an exclusion criterion but patients refrained from these drugs 48 hours before sessions. Participants with MRI contraindications (e.g. metal implant) were included but not enrolled in MRI. (Table 1, Supplementary Figure 1 for CONSORT diagram).

**Table 1.**
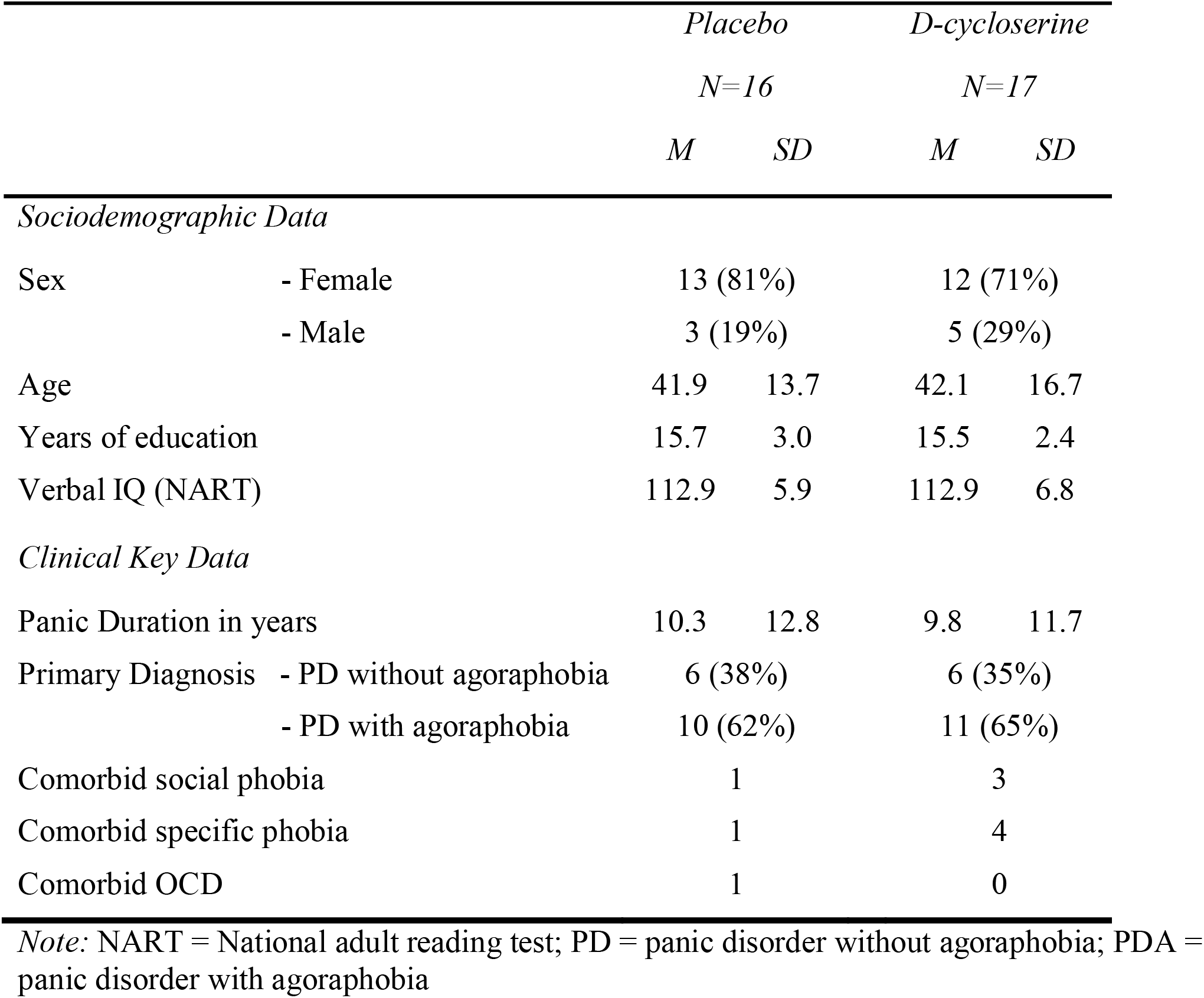
Demographic and Clinical Key Data in placebo versus d-cycloserine group.

### General Procedure

Figure 1 gives an overview of assessments. Participants were randomly allocated to a single oral dose of d-cycloserine (250 mg; King Pharmaceuticals) or placebo capsule (microcrystalline cellulose) (1:1 ratio), administered 2 h before single-session CBT. Previous work into the effects of d-cycloserine on CBT often used doses of 50mg, but different from our study these trials used longer CBT protocols, and healthy volunteer work indicates an effect of d-cycloserine on single-session hippocampal learning at 250mg but not 50mg (7). Considering that we only offered one session, and based on work showing no significant differences in clinical effects of ultra-brief CBT together with d-cycloserine between 50mg and 500mg, we chose a dose of 250mg (8).

**Figure 1.**
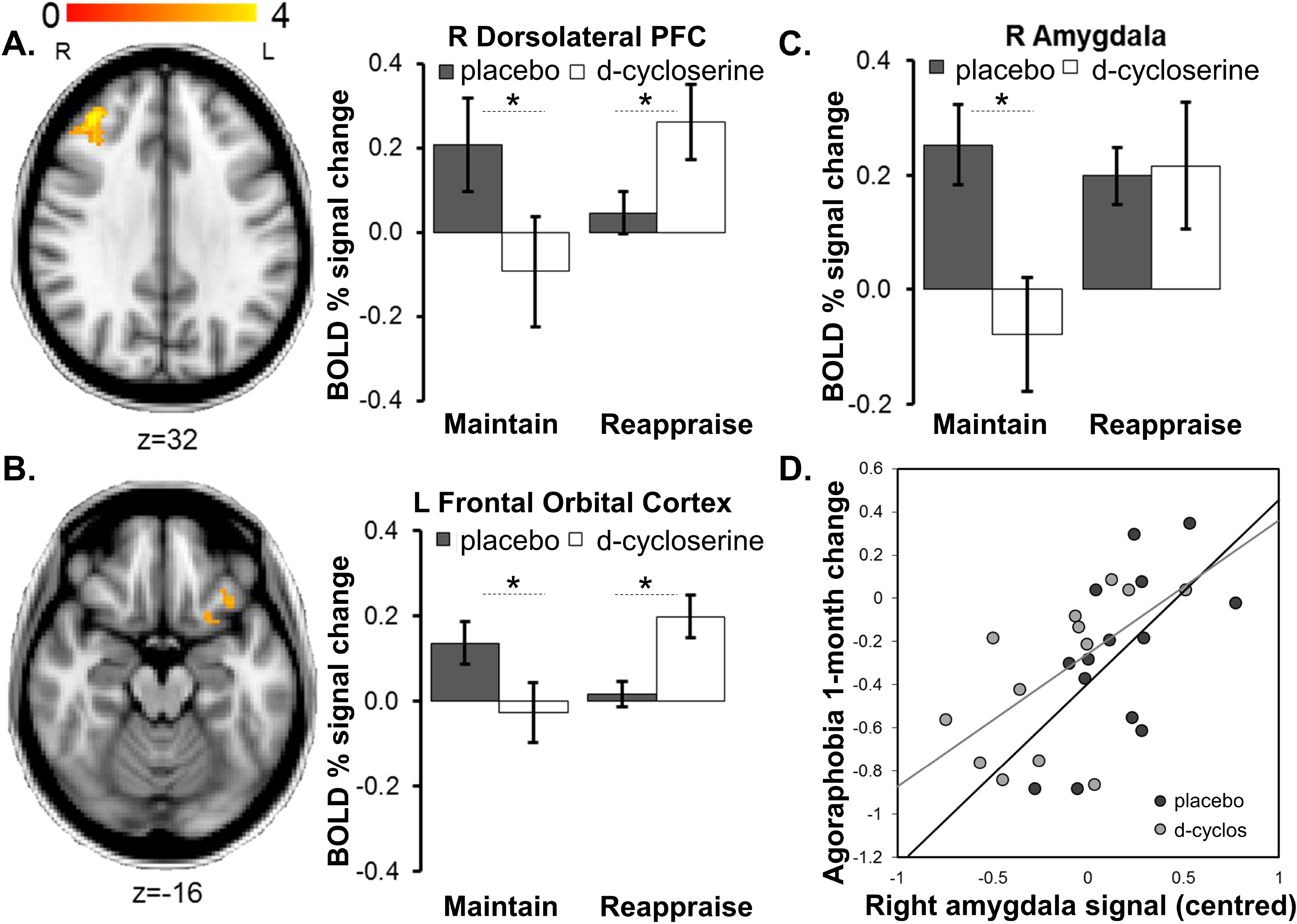
Overview of study visits and assessments throughout the trial.

All sessions took place in a university setting. After screening, participants returned for four study visits, with baseline assessment and intervention taking place on day 1, and outcome testing taking place 1 day, 1 month, and 6 months after intervention. During the screening, sociodemographic (age, gender, years of education, verbal intelligence) and basic clinical data (primary panic diagnosis, comorbidity, panic duration) were acquired (9). Clinical symptoms were assessed on all four visits. On the day after intervention, we also assessed emotional processing using behavioural computer tasks and magnetic resonance imaging (MRI). Recruitment, obtaining consent, eligibility screening, sending enrolment requests to pharmacy, double-blind drug administration, exposure therapy and clinical and neurocognitive testing were carried out by a trained clinical psychologist researcher (AR). All procedures were in accordance with the Declaration of Helsinki and approved by a research ethics committee. Participants gave written informed consent. Data were collected between November 2013 and April 2016.

### Single-session Exposure Therapy

Treatment was delivered by an experienced clinical psychologist. The intervention followed a previously published protocol and was a condensed version of routine clinical care (3, 10). It involved one session of exposure-based CBT, based on the well-established cognitive-behavioural theory of panic (11). Session ingredients: (i) cognitive preparation: explanation of individually relevant learning mechanisms underlying the maintenance of anxiety, especially the role of safety strategies (~15min), (ii) exposure to a fear-provoking situation (e.g. being locked in walk-in closet) and bodily sensations while dropping all safety strategies, to test out catastrophic expectations and break through stimulus-driven response cycles (15min exactly for each participant), (iii) cognitive debriefing, to discuss the patient’s experience in a fear situation without safety strategies, and to consolidate new behaviour (~10min).

### Randomisation and Masking

Participants and the researcher responsible for treatment, data collection and outcome evaluation remained naïve to drug group allocation until completion of data analysis. Placebo was encapsulated in lactose capsules identical to d-cycloserine. Generation of the randomisation sequence, treatment allocation and drug dispensing were executed by an external pharmacy not in direct contact with study participants, and the allocation list remained in pharmacy until all steps of data blind-analysis had been completed. The randomization sequence was generated using a random number generator (www.random.org) and was based on blocked randomization (blocks of four) while stratifying for gender and primary diagnosis (panic disorder either with or without agoraphobia).

### Outcomes

#### Primary Outcomes

Primary outcome measure was a faces dot probe task (FDOT). Stimuli were coloured photographs of 20 individual faces with a neutral, fearful and happy facial expression each (12). In each of 192 trials, a neutral-neutral, neutral-happy, or neutral-fearful face pair was presented, with one stimulus appearing above and one below a central fixation position. Eight blocks of unmasked versus masked trials each were presented alternatingly. Faces pairs were presented for 100ms in the unmasked and 16ms (plus scrambled face pair for 84ms) in the masked condition, followed by a dot probe that participants categorised as horizontal or vertical using two buttons. Position of an emotional face, probe position and type were fully counterbalanced. This task design involved congruent trials (dot at the position of an emotional face) and incongruent trials (dot replaces neutral face while emotional face is present). An implicit association task (13), a second primary outcome, lead to no significant group effects and is therefore not described here.

#### Secondary Outcomes

##### i) MRI

Patients performed an emotion regulation task on the day after treatment during 3T MRI (14). IAPS images showing negatively valenced panic-related scenes (e.g. funeral) were presented in 8 blocks of 5 images (5s each), alternating with grey fixation baseline blocks (30s) (15). For half of the blocks, participants were instructed to naturally experience the emotional state evoked by the images, without attempting to regulate it (Maintain blocks). For Reappraisal blocks, they were instructed to down-regulate the provoked negative affect by using previously demonstrated strategies of cognitive reappraisal (e.g. reframing, rationalising). Individual contrast images were calculated for Maintain blocks, Reappraisal blocks, Maintain versus Reappraisal and Reappraisal versus Maintain. Based on our previous work which identified the amygdala as hyperactive in Maintain but not Reappraisal blocks in patients with panic disorder, we ran region of interest analyses (ROI), including 10mm radius spherical masks around the peak voxel of a left amygdala region (−14/−6/−8) and its right-hemisphere counterpart, with Maintain BOLD % signal as the main outcome measure. We further explored group effects running a mixed-effects whole-brain analysis ((14) for image acquisition and analysis details).

##### ii) Clinical Symptom Measures

Before treatment, on the day after, and at 1-month and 6-month follow-up, anxiety and depression symptoms were monitored using established self-report measures: i) State-Trait Anxiety Inventory (STAIS/STAIT), ii) Beck Depression Inventory (BDI), iii) Body Sensations Questionnaire assessing fear of physical sensations (BSQ), iv) Agoraphobic Cognitions Questionnaire (ACQ), v) Mobility Inventory (MI) assessing agoraphobic avoidance, and vi) Panic Attack Scale (PAS) assessing panic frequency (16–20). The clinician-administered Panic Disorder Severity Scale (PDSS) was used at baseline and follow-up appointments (21). To establish in-vivo stress reactivity, anxiety was also measured during 5min exposure to an individually relevant agoraphobic situation, before and after the day of treatment and at follow-ups (3, 10). This involved being locked in an enclosed walk-in closet (17 CYC/14 PLAC), or a bus or car drive (0 CYC/2 PLAC). Immediately after the test, participants rated their level of anxiety experienced during the test (before, after 1min, after 3min, end of test), using 0-100 visual analogue scales. Baseline ratings were acquired during an earlier part of the sessions.

### Demand and Side Effects and Concomitant Treatment

To capture any acute changes, blood pressure and heart rate were measured before and 2hrs after drug administration (expected peak level), and participants completed visual analogue scales rating their mood and physiological symptoms. At the end of the intervention day, participants and experimenter also guessed whether the active capsule had been administered.

### Statistical Analysis

#### Sample Size

We predicted that the placebo group would show next-day fear bias similar to that seen after single-session CBT (*M*=5, *SD*=25), and that the d-cycloserine group would show next-day fear bias similar to that seen in healthy volunteers (*M*=−15, *SD*=25) in our previous work (3). With an effect size of d=0.8 and an α-level of 0.05, 16 participants per group would be needed to achieve 70% power. We originally aimed to recruit 50 patients for this study but this was adjusted to 32 based on the above.

#### Overall Statistical Approach

Data analyses included all randomised subjects and were performed on the intention-to-treat basis (16 PLAC, 17 CYC), except for MRI measures which were analysed per protocol (14 PLAC, 13 CYC). No data was excluded from analysis. To account for participants lost to follow-up (Supplementary Figure 1), missing data were imputed, assuming data missing at random, by means of multiple imputation by chained equations (incomplete cases: 1-month follow-up 6% PLAC/ 6% CYC, 6-months follow-up: 19% PLAC/ 18% CYC). Twenty complete data sets were generated using the statistical computing software R (R Core Team, 2018), with 20 iterations per imputation (22). Predictors were sociodemographic and clinical baseline characteristics, and baseline and next-day clinical outcome measures; imputed variables were clinical outcomes at follow-ups. Statistical analyses were run in R using the packages nlme (for linear mixed effects models) and miceadds (additional statistics for multiple imputation). Repeated measures from the same participant were accounted for by means of random effects. In all analyses, the stratification variables gender and primary diagnosis were used as covariates. Statistical tests were two-tailed and based on an alpha-level of significance of 0.05. Treatment effect sizes are reported as Cohen’s *d*.

#### Cognitive Biomarker Measures

In the FDOT, reaction times (RT) below 200 ms and above 2000 ms, or above 2 SDs above the mean, were excluded (3, 23). Bias scores were calculated by subtracting the median RT in congruent trials from the median RT in incongruent trials. Group differences were assessed using linear mixed-effects models, including the fixed factors group and condition (happy, fear). MRI data were analysed using FSL FEAT 6.0 (FMRIB Software Library; www.fmrib.ox.ac.ul/fsl) with Z>2.3 and *p*<0.05. Extracted ROI BOLD % signal changes were entered into linear mixed-effects models with the factors group and condition (Maintain, Reappraisal).

#### Clinical Symptom Measures

Group differences on the validated clinical outcome measures and stress test ratings were assessed using linear mixed-effects models, including the factors group and time (next day, 1-month, 6-months) and their interaction, and controlling for pre-treatment scores. In line with recent work, recovery was defined as agoraphobia scores (MI) falling within the range reported for healthy subjects, as this measure most reliably reflects the impact of the disorder on daily functioning (3, 16). Following established approaches, we also ran eight separate multiple linear regression analyses to predict whether i) BOLD % amygdala signal change (Maintain block) and ii) threat bias (FDOT) on the day after treatment predicted symptom changes at 1-month follow-up on the MI, BSQ, ACQ, or PDSS (3, 24). Next-day symptom severity (baseline for PDSS) on that measure was entered as predictor of no interest to control for its potential influence on the outcome at 1 month. Group, bias, and the group-bias interaction term were entered as predictors of interest.

## Results

### Primary Outcomes

Next-day threat bias for fearful faces was significantly lower in the d-cycloserine compared to the placebo group (*p*=0.042, *d*=0.77) (Table 2).

**Table 2.** Primary and secondary outcomes at baseline, on the day after treatment, and at 1-month and 6-months follow-ups.

|  |  | PLACEBO | D-CYCLOSERINE | Adjusted mean diff (SE) | 95% CI, p-value, Cohen's d |
| --- | --- | --- | --- | --- | --- |
| **FDOT (fear)** | Next-Day | 21.3 (30.8) | 2.1 (17.2) | -19.0 (9.3) | -38.04 to 0.14; **p=0.042**; d=0.77 |
| **Right Amygdala** | Next-Day | 0.25 (0.27) | -0.08 (0.35) | -0.32 (0.14) | -0.61 to -0.03; **p=0.023**; d=1.06 |
| **Left Amygdala** | Next-Day | 0.17 (0.33) | 0.05 (0.68) | -0.12 (0.19) | -0.50 to 0.27; p=0.54; d=0.22 |
| **STAI-S (20-80)** | Baseline | 42.8 (7.3) | 43.9 (12.5) | .. | .. |
|  | Next-Day | 44.9 (8.8) | 34.6 (11.0) | -10.67 (3.57) | -17.83 to -3.51; **p=0.0041**; d=1.01 |
|  | 1-month | 39.7 (9.2) | 32.2 (10.9) | -7.92 (3.71) | -15.30 to -0.54; **p=0.036**; d=0.74 |
|  | 6-months | 38.9 (12.8) | 31.4 (9.7) | -8.05 (3.95) | -15.94 to -0.16; **p=.046**; d=0.66 |
| **STAI-T (20-80)** | Baseline | 52.2 (6.7) | 52.4 (9.4) | .. | .. |
|  | Next-Day | 50.8 (8.3) | 47.1 (10.7) | -3.53 (3.60) | -10.74 to 3.68; p=0.33; d=0.39 |
|  | 1-month | 46.3 (11.8) | 38.2 (12.6) | -7.99 (3.74) | -15.44 to -0.54; **p=0.036**; d=0.66 |
|  | 6-months | 44.4 (12.8) | 35.2 (10.3) | -9.20 (3.89) | -16.96 to -1.44; **p=0.021**; d=0.79 |
| **BDI (0-63)** | Baseline | 16.8 (9.4) | 17.3 (11.7) | .. | .. |
|  | Next-Day | 18.3 (10.1) | 13.2 (10.0) | -5.26 (2.46) | -10.19 to -0.33; **p=0.037**; d=0.51 |
|  | 1-month | 12.5 (8.0) | 11.1 (10.4) | -1.57 (2.56) | -6.66 to 3.53; p=0.54; d=0.15 |
|  | 6-months | 12.0 (10.1) | 6.3 (6.6) | -5.86 (2.72) | -11.29 to -0.42; **p=0.035**; d=0.67 |
| **ACQ (1.0-5.0)** | Baseline | 2.5 (0.5) | 2.4 (0.7) | .. | .. |
|  | Next-Day | 2.6 (0.5) | 2.3 (0.6) | -0.21 (0.19) | -0.60 to 0.17; p=0.27; d=0.30 |
|  | 1-month | 2.2 (0.7) | 1.9 (0.7) | -0.29 (0.20) | -0.69 to 0.11; p=0.16; d=0.43 |
|  | 6-months | 2.0 (0.7) | 1.6 (0.6) | -0.37 (0.22) | -0.81 to 0.07; p=0.095; d=0.61 |
| **BSQ (1.0-5.0)** | Baseline | 3.2 (0.8) | 3.2 (0.7) | .. | .. |
|  | Next-Day | 3.2 (0.8) | 3.0 (0.8) | -0.12 (0.27) | -0.66 to 0.41; p=0.65; d=0.14 |
|  | 1-month | 2.6 (0.8) | 2.5 (0.8) | -0.06 (0.27) | -0.60 to 0.48; p=0.82; d=0.13 |
|  | 6-months | 2.4 (1.0) | 2.2 (1.0) | -0.12 (0.29) | -0.71 to 0.47; p=0.68; d=0.20 |
| **MI (1.0-5.0)** | Baseline | 2.3 (0.7) | 2.1 (0.9) | .. | .. |
|  | Next-Day | 2.2 (0.5) | 2.1 (0.8) | -0.038 (0.16) | -0.35 to 0.27; p=0.81; d=0.15 |
|  | 1-month | 1.9 (0.6) | 1.7 (0.8) | -0.12 (0.16) | -0.43 to 0.20; p=0.46; d=0.28 |
|  | 6-months | 1.8 (0.5) | 1.5 (0.6) | -0.18 (0.18) | -0.54 to 0.19; p=0.34; d=0.54 |
| **PAS (0-4)** | Baseline | 1.8 (1.1) | 1.7 (1.2) | .. | .. |
|  | Next-Day | 1.8 (1.0) | 1.5 (1.0) | -0.26 (0.27) | -0.79 to 0.27; p=0.33; d=0.30 |
|  | 1-month | 0.7 (0.8) | 0.5 (1.0) | -0.22 (0.30) | -0.81 to 0.37; p=0.46; d=0.22 |
|  | 6-months | 0.5 (0.8) | 0.3 (0.7) | -0.19 (0.30) | -0.79 to 0.40; p=0.52; d=0.27 |
| **PDSS (0-28)** | Baseline | 11.8 (4.0) | 11.6 (4.6) | .. | .. |
|  | 1-month | 6.1 (4.4) | 4.8 (5.3) | -1.31 (1.39) | -4.11 to 1.48, p=0.35, d=0.27 |
|  | 6-months | 4.6 (3.8) | 3.4 (3.8) | -1.19 (1.45) | -4.12 to 1.73, p=0.41; d=0.24 |
| **Stress (0-100)** | Baseline | 41.1 (25.9) | 43.4 (24.7) | .. | .. |
|  | Next-Day | 29.2 (21.5) | 17.8 (14.6) | -14.09 (4.55) | -23.0 to -5.1; **p=0.0021**; d=0.62 |
|  | 1-month | 19.1 (17.3) | 13.0 (15.2) | -8.84 (4.63) | -17.9 to 0.3; p=0.057; d=0.37 |
|  | 6-months | 20.0 (22.0) | 11.5 (15.6) | -11.20 (4.67) | -20.4 -2.0; **p=0.017**; d=0.45 |

*Note:* FDOT = Faces Dot Probe Task; IAT = Implicit Association Task; STAI-S/ STAI-T = state trait anxiety inventory; BDI = Beck Depression Inventory; ACQ = Agoraphobic Cognitions Questionnaire; BSQ = Body Sensations Questionnaire; MI = Mobility Inventory; PAS = Panic Attack Scale; PDSS = Panic Disorder Severity Scale.
STAIS
STAIT
BDI
ACQ
BSQ
PAS
PDSS

### Secondary Outcomes

#### MRI

D-cycloserine versus placebo lead to a significant decrease of right amygdala activation in Maintain (*p*=0.023) but not Reappraisal blocks (mean difference 0.02, 95% CI −0.23 to 0.27; *p*=0.88). These effects are in line with previous work showing increased threat response during Maintain but not Reappraisal blocks in patients compared to controls (14). No drug effects were found in the left amygdala (Maintain: *p*=0.54; Reappraisal: mean difference −0.08, 95% CI −0.47 to 0.30; *p*=0.66) (Table 2; Figure 2 C). A whole-brain analysis identified significant group by task effects in the left frontal orbital (OFC; 499 voxels, MNI −28,12,−14, Z=3.40, p=0.003) and right dorsolateral prefrontal cortex (DLPFC; 379 voxels, MNI 36,36,30, Z=4.06, p=0.017), suggesting diminished activation in these regulatory areas in Maintain blocks after d-cycloserine (Figure 2 A/B).

**Figure 2.**
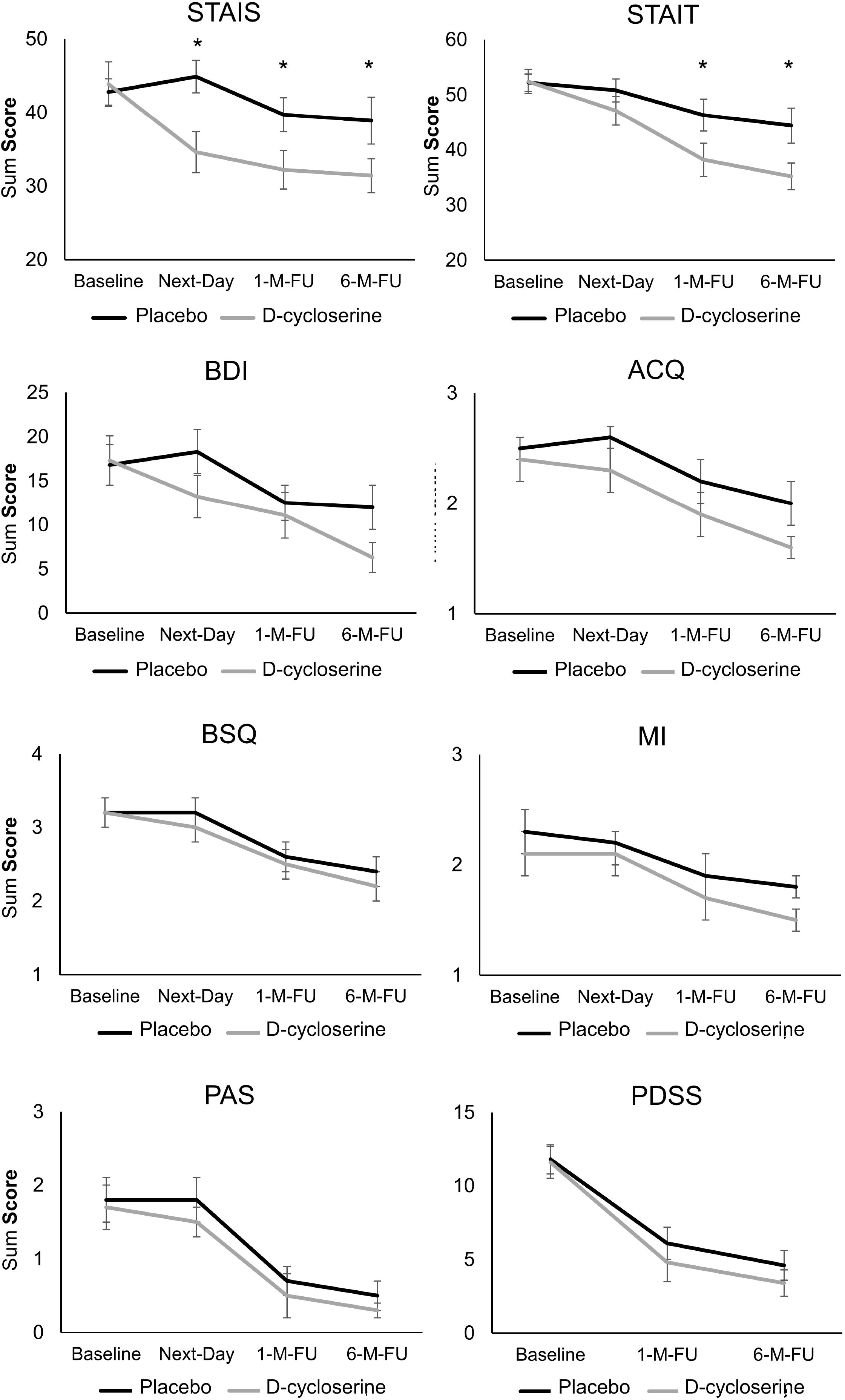
**A./B.** Whole-brain analysis Group x Task interaction: Compared to placebo, d-cycloserine-augmented single-session CBT led to significantly stronger activation in areas of cognitive control during Reappraise versus Maintain on the day after treatment. **C.** Region-of-interest analysis Group x Task interaction: Compared to placebo, cycloserine lead to reduced activation of the right amygdala during Maintain block. **D.** The relationship between amygdala responsivity on the day after single-session CBT and symptom improvement at 1-month follow-up: lower amygdala responsivity on the day after treatment predicts stronger reduction in agoraphobia severity as measured using the self-report Mobility Inventory (MI) across groups. Images thresholded at *Z*>2.3, *p*<0.05, corrected. Note: Error bars show SEM. An asterisk indicates an alpha-level of significance p<0.05.

#### Clinical Symptom Measures

There were no significant group differences in fear of physical symptoms (BSQ), agoraphobia severity (MI), agoraphobic cognitions (ACQ), panic attack frequency (PAS) or panic severity (PDSS) on the day after treatment or at follow-ups (all *p*>0.095) (Table 2; Figure 3). On the day after treatment, the d-cycloserine compared to placebo group reported lower state anxiety (STAIS; *p*=0.0041) and depression scores (BDI *p*=0.0037). The STAIS group difference remained sustained throughout follow-ups (both *p*<0.0046); the BDI group difference reduced at 1-month follow-up due to the placebo group achieving additional gains (*p*=0.54) but reappeared at 6-month follow-up (*p*=0.035) due to the d-cycloserine group showing further response. Trait anxiety was significantly lower after d-cycloserine-versus placebo-supported CBT at 1- and 6-months follow-ups (both *p*<0.0036). In the stress test, the d-cycloserine group reported lower anxiety on the day after treatment and at 6-months follow-up (both *p*<0.0021), but the group difference failed to reach statistical significance 1-month after treatment (*p*=0.057).

**Figure 3.**
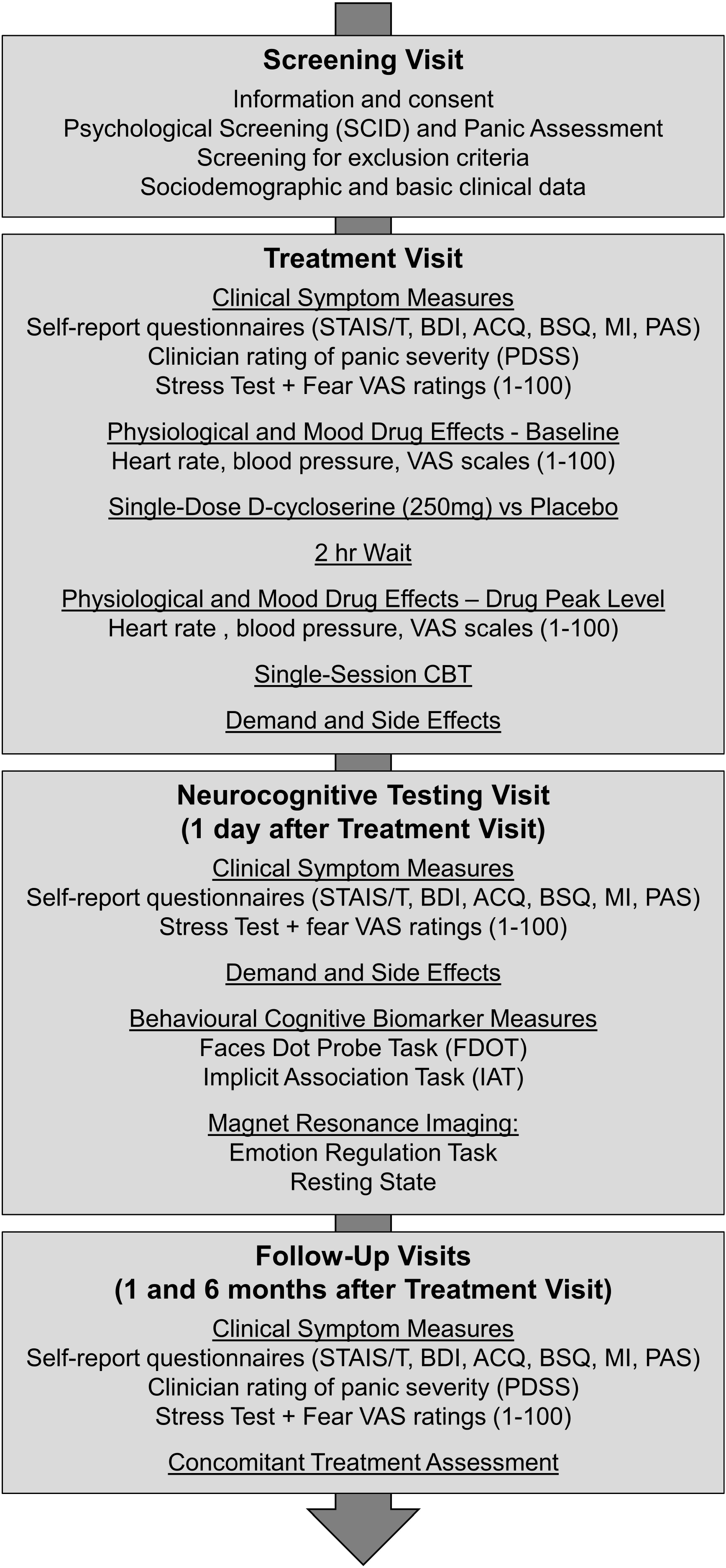
Self-report and clinician-rated symptom scores for the two groups at baseline, on the day after treatment and at follow-ups, after multiple imputation of missing data (error bars show SEM). D-cycloserine lead to stronger improvements in state and trait anxiety (STAIS, STAIT), depressivity (BDI) and agoraphobic cognitions (ACQ) on the day after treatment, with group differences remaining significant throughout follow-ups. Note: Error bars show SEM. An asterisk indicates an alpha-level of significance p<0.05.

Recovery rates were still low and similar between groups on the day after treatment (PLAC 4/16 (25%), CYC 5/17 (29.4%), *RR* 1.06, *CI* 0.70-1.61, *p*=0.98). At 1-month follow-up, the d-cycloserine group showed significantly greater treatment response than the placebo group (PLAC 4/16 (25.0%), CYC 12/17 (70.1%), *RR* 2.55, *CI* 1.16-5.61, *p*=0.015). At 6-months follow-up, the placebo group showed improved clinical gains, closer to those of the d-cycloserine group (PLAC 7/16 (43.8.0%), CYC 12/17 (70.6%), *RR* 1.91, *CI* 0.81-4.49, *p*=0.17).

### Prediction of Clinical Symptom Changes

FDOT threat bias was not predictive of clinical change (all *R^2^*<0.20, all *p*>0.25). Amygdala responsivity on the day after treatment predicted 1-month follow-up changes in agoraphobia severity (MI) across groups (*R^2^*=0.48; main effect bias: *p*=0.0070; other outcomes: all *R^2^*<0.38, all *p*>0.11), with patients showing the lowest amygdala response achieving greater reduction in avoidance during follow-up (Figure 2D).

### Demand and Side Effects and Concomitant Treatment

No serious adverse events were reported. D-cycloserine caused no acute differential changes in blood pressure, heart rate and mood (Supplementary Table 1). Neither experimenter nor participants were able to correctly guess group allocation (d-cycloserine guesses; experimenter PLAC 44%, CYC 41%, *p*=0.88; participant PLAC 25%, CYC 53% *p*=0.10), suggesting that double-blindness was maintained.

## Discussion

Threat bias for fearful faces (FDOT) and amygdala response to aversive images were significantly lower in the d-cycloserine compared to the placebo group on the day after treatment. Greater next-day reduction in amygdala threat response was associated with greater improvement in agoraphobic avoidance during 1-month follow-up. D-cycloserine led to significantly greater clinical recovery at 1-month follow-up than placebo (71% vs 25%), but recovery rates were not statistically different at 6-months follow-up (71% vs 44%). There were no group differences on clinical measures specific to panic disorder, but the drug improved response to an in-vivo stress test and outcome on more global measures of mental health, including state-trait anxiety and depression, with medium to large effects.

These findings highlight a possible neuropsychological mechanism of action of exposure-based CBT that might also be a crucial interface for d-cycloserine action. Across participants, lower amygdala response on the day after treatment predicted lower symptom severity one month later, suggesting that this neural effect might be a key mechanism by which exposure therapy exerts its clinical effects. The amygdala is thought to be crucial in threat processing, and increased responsivity is characteristic of anxiety disorders (25). In contrast, a reduction in amygdala hypersensitivity is seen after exposure therapy, where threat stimuli lose their potential to automatically signal danger (25, 26). Our results suggest that this change in amygdala function occurs very rapidly during CBT, after only one session. Over time and in interaction with everyday challenges, this reduced threat sensitivity presumably translates into more distinct symptom improvement. These findings add to our previous work showing that attentional bias for fearful faces mediates clinical outcome (3).

Even though this study identified no specific neuropsychological mechanism of d-cycloserine action, the drug significantly reduced attention bias and amygdala threat responsivity within one day of treatment. These findings corroborate the idea that threat processing might be a landmark parameter for treatment enhancement with d-cycloserine. The drug is thought to enhance NMDA receptor functioning and neuroplasticity in the amygdala and hippocampus, areas relevant to fear extinction and threat processing (27). It is possible that d-cycloserine exerts its augmentative clinical effects by further amplifying changes in amygdala sensitivity taking place during exposure. As our results show, these changes occur early during treatment, providing a rationale as to why d-cycloserine has particularly beneficial effects on clinical outcome when applied early in the therapeutic process.

Although our clinical results remain to be replicated in a large-scale clinical trial, this study also replicates earlier findings showing that 25% of patients reach recovery one month after single-session CBT (3). The present results add to this observation, indicating that these clinical effects are not only stable during a follow-up of 6 months, but that additional clinical gains may be achieved during this time-frame. This study also provides preliminary evidence that d-cycloserine may improve clinical response to single-session CBT. 71% of participants in the d-cycloserine group fulfilled criteria for recovery at 1- and 6-month follow-up, a rate comparable to standard longer-term CBT courses (28). These findings point to the possibility that if used in an optimal way, targeting very early therapeutic learning, d-cycloserine may lead to substantially improved clinical effects.

While these results are promising for the development of more compact psychological-pharmacological combination treatments, there are limitations. While we previously found that FDOT threat bias one day after single-session CBT predicted clinical improvement, this observation was not replicated in this study. Recent work into the psychometric properties of the dot probe task suggests weak test-retest reliability, indicating that alternative measures of threat bias might be preferable (29). In line with this recommendation, we found that amygdala threat response was more sensitive to brief treatment than the behavioural measures used, leading to large effect sizes of d>1 and predicting clinical recovery. However, MRI and threat bias measures were only applied after but not before treatment, limiting our ability to evaluate to what degree observed between-group differences on the day after treatment relate to within-subject change. Furthermore, this study was designed to detect neurocognitive rather than clinical effects of d-cycloserine, and even though our clinical observations indicate efficaciousness of single-session CBT with d-cycloserine, these findings need to be validated in a large-scale trial to reliably evaluate clinical efficacy of this combination intervention. While our participants are representative of those seen in routine clinical care regarding symptom severity and sociodemographic markers, they were recruited from the community rather than being treatment-seekers, and they were unmedicated (30). Future trials will have to investigate whether the facilitative effects of d-cycloserine on single-session CBT found here translate to these cases.

Taken together, this is the first study to identify a neurocognitive mechanism by which d-cycloserine may deploy its augmentative effects on clinical outcomes during exposure therapy for anxiety. The drug substantially amplifies changes in amygdala threat reactivity taking place early in psychological treatment, and the magnitude of these changes predicts clinical improvement across time. Our findings suggest that threat processing might be a landmark parameter for treatment enhancement with d-cycloserine, and they provide a possible explanation as to why d-cycloserine particularly affects clinical outcome when applied early in treatment.

## Supporting information

SuppFig1

SuppTab1

## Acknowledgements

This research was funded by a Medical Research Council Experimental Medicine grant awarded to CH and AR (MR/J011878/1) and by a John Fell Oxford University Press (OUP) research grant awarded to AR (123/807), and it was supported by the NIHR Oxford Health Biomedical Research Centre. AR is supported by a fellowship from MQ: Transforming Mental Health. MB is supported by a fellowship from the MRC. The funders had no role in the design of the study, data collection or interpretation of results.

## Notes

#### Summary of Updates

shortened in overall length without changing overall content or interpretation

