## Supplementary figures and images for "Neurocognitive mechanisms of d-cycloserine augmented single-session exposure therapy for anxiety"

### SuppFig1

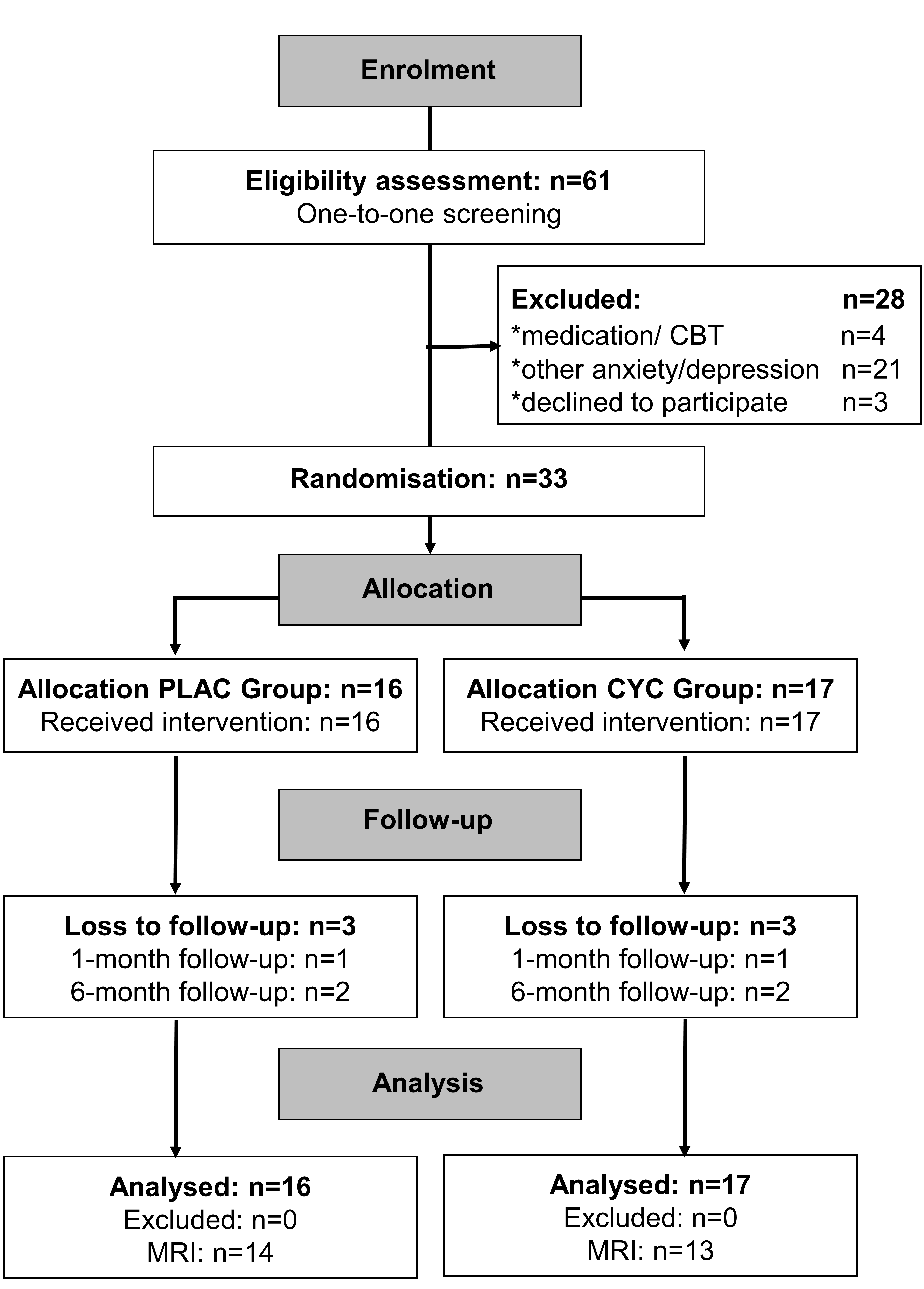
