## Supplementary material for "Neurocognitive mechanisms of d-cycloserine augmented single-session exposure therapy for anxiety": SuppTab1

**Supplementary Table 1. Physiological measures and visual analogue scale ratings in the two groups before drug intake and post drug (M, SD).** Cycloserine compared to placebo caused no acute differential changes in blood pressure, heart rate or mood: Time (baseline, drug peak) by Group repeated measures ANOVAs run for heart rate, blood pressure scores, and each of the VAS items indicated no significant main effects of group and no significant Group x Time interaction effects for blood pressure, heart rate, and most of the VAS items (all F(1,31)<2.5, all p>0.13). For the item ‘nauseous’, there was a significant Group x Time interaction (F(1,31)=5.4, p=0.027), driven by the placebo group feeling less nauseous at drug peak level compared to baseline (cycloserine: t(16)=0.4, p=0.68; placebo: t(15)=3.1, p=0.007; between-group t-tests for baseline and drug peak both t(31)<1.35, both p>0.19).

|  | **Placebo**  **N=16** | |  | **D-Cycloserine**  **N=17** | |
| --- | --- | --- | --- | --- | --- |
|  | **M** | **SD** |  | **M** | **SD** |
| **BASELINE** |  |  |  |  |  |
| **Physiological Measures** |  |  |  |  |  |
| Heart rate | 70·5 | 10·0 |  | 69·0 | 11·5 |
| Blood pressure - systolic | 124·9 | 19·5 |  | 132·5 | 29·6 |
| Blood pressure – diastolic | 80·3 | 13·2 |  | 81·7 | 20·5 |
| **Visual Analogue Ratings VAS** |  |  |  |  |  |
| Anxious | 43·3 | 19·0 |  | 36·6 | 22·6 |
| Tearful | 7·3 | 9·2 |  | 6·7 | 10·8 |
| Hopeless | 19·4 | 18·4 |  | 14·5 | 21·6 |
| Sad | 19·2 | 14·4 |  | 15·9 | 23·0 |
| Depressed | 21·2 | 21·2 |  | 17·3 | 18·5 |
| Sleepy | 42·1 | 23·8 |  | 41·7 | 27·3 |
| Nauseous | 20·2 | 21·2 |  | 10·9 | 18·1 |
| Dizzy | 11·9 | 10·9 |  | 15·4 | 22·3 |
| Heart racing | 6·7 | 5·9 |  | 14·1 | 20·3 |
| Alert | 52·6 | 17·2 |  | 47·5 | 21·6 |
| **POST DRUG** |  |  |  |  |  |
| **Physiological Measures** |  |  |  |  |  |
| Heart rate | 70·9 | 9·5 |  | 64·3 | 9·2 |
| Blood pressure - systolic | 125·0 | 18·6 |  | 135·9 | 32·8 |
| Blood pressure – diastolic | 77·2 | 8·5 |  | 82·8 | 20·6 |
| **Visual Analogue Ratings VAS** |  |  |  |  |  |
| Anxious | 18·1 | 11·2 |  | 13·2 | 16·6 |
| Tearful | 5·1 | 5·5 |  | 4·2 | 8·7 |
| Hopeless | 6·3 | 6·3 |  | 8·7 | 13·9 |
| Sad | 8·0 | 14·0 |  | 8·0 | 14·0 |
| Depressed | 12·6 | 14·2 |  | 8·4 | 15·1 |
| Sleepy | 46·3 | 27·3 |  | 35·6 | 30·4 |
| Nauseous | 8·9 | 8·2 |  | 9·9 | 14·0 |
| Dizzy | 7·3 | 7·5 |  | 14·9 | 19·6 |
| Heart racing | 7·3 | 9·2 |  | 11·5 | 16·1 |
| Alert | 48·8 | 21·0 |  | 48·1 | 22·0 |
